# Hedonic hotspot in rat olfactory tubercle: map for mu-opioid, orexin, and muscimol enhancement of sucrose ‘liking’

**DOI:** 10.1101/2025.11.18.689156

**Authors:** Koshi Murata, Kent C. Berridge

## Abstract

Pleasure plays a crucial role in positive reinforcement and motivation. Brain regions able to amplify positive hedonic reactions to sweetness, known as ‘hedonic hotspots’, are distributed within the mesocorticolimbic reward systems. The olfactory tubercle (OT), a part of the ventral striatum that receives olfactory input, contains distinct functional domains: the anteromedial domain mediates approach motivation toward odors associated with food, whereas the lateral domain mediates avoidance motivation away from odors associated with danger. However, it has remained unclear whether the OT modulates hedonic reactions to pleasant sensations. In this study, we made pharmacological microinjections in OT of in rats to examine whether these OT subregions can modulate hedonic reactions, as assessed by the taste reactivity test. Sweet oral infusions of sucrose solution were delivered into the mouth via an intraoral cannula, and the rats’ orofacial and somatic hedonic reactions were recorded and analyzed. We compared three pharmacological agents: mu-opioid receptor agonist DAMGO, orexin-A peptide, and GABA_A_ receptor agonist muscimol. Microinjection of any of these drugs into the anteromedial OT subregion enhanced hedonic ‘liking’ reactions to sucrose. Furthermore, DAMGO injection into the anteromedial OT subregion recruited distant Fos expression in other ‘hedonic hotspots’, including in the caudal ventral pallidum and the rostromedial orbitofrontal cortex. By contrast, the same microinjections into the anterolateral OT subregion failed to enhance ‘liking’ reactions and, DAMGO oppositely increased aversive ‘disgust’ reactions. These findings suggest that the anteromedial OT contains a ‘hedonic hotspot’, whereas the anterolateral OT may contain a suppressive opioid ‘hedonic coldspot’. Thus, OT subregions may help causally modulate hedonic reactions to sweetness and flavor perception.

## Introduction

Hedonic reactions to pleasant sensations play roles in motivating adaptive behaviors of animals and humans essential for survival. To elucidate neurobiological mechanisms underlying pleasure, it is essential to objectively measure affective ‘liking’ reactions elicited by the hedonic impact of pleasant stimuli such as sensory rewards. Affective neuroscience studies have often used the taste reactivity test, which quantifies distinct orofacial expressions evoked by pleasant versus unpleasant tastes, to measure ‘liking’ reactions to sweet tastes and ‘disgust’ reactions to bitter tastes^1,2^. Rodents, non-human primates, and human infants share many of these affective facial reactions.

Pharmacological microinjections and optogenetic manipulations of neural systems have revealed hedonic mechanisms by mapping the ability of neurobiological modulations to causally change hedonic orofacial ‘liking’ reactions to tastes^3,4^. In particular, this mapping has identified multiple hedonic hotspots across the brain, which are small restricted subregions within mesocorticolimbic structures that are especially able to enhance ‘liking’ reactions to sweetness in response to local neurobiological manipulations. Identified hedonic hotspots include the rostrodorsal quadrant of medial shell of nucleus accumbens (NAc), the caudolateral half of ventral pallidum (VP), a rostromedial portion of the orbitofrontal cortex (OFC), a far posterior zone of the insula cortex, and the parabrachial nucleus of the brainstem pons^5–9^. These hedonic hotspots that generate ‘liking’ are embedded within larger mesocorticolimbic circuitry that can generate incentive salience or ‘wanting’, suggesting a close interconnection between ‘liking’ and ‘wanting’ mechanisms in reward^10^.

The olfactory tubercle (OT), despite its olfactory name, is actually a rostroventral extension of the ventral striatum, sharing several neurobiological features and being anatomically continuous with the NAc^11–13^. For example, both the OT and NAc contain GABAergic principal neurons known as medium spiny neurons (MSNs), whose primary axonal target is the VP. Both OT and NAc MSNs express dopamine receptors, mostly expressing either D1 dopamine receptors and the opioid peptide dynorphin, or D2 dopamine receptors and the opioid peptide enkephalin^14,15^. Therefore, the OT has also been referred to as the “olfactory striatum” or “tubular striatum”^16,17^.

The OT also mediates reward and motivation functions similarly to NAc. For example, rats self-administer intracranial microinjections of addictive drugs in OT, particularly within its anteromedial subregion (OTam)^18–20^. Involvement of OTam in odor-guided appetitive behaviors has been demonstrated by mapping Fos activation elicited during attraction to an odor cue paired with sugar-reward^21^. Further, mesolimbic dopamine inputs from the ventral tegmental area (VTA) to the medial OT mediate the acquisition and execution of odor preference^22^. By contrast, the anterolateral OT (OTal) appears to participate in aversive behavior, as indicated by Fos activation in OTal during aversion to an odor cue paired with foot-shock^21^. Similarly, the caudal subregion of NAc medial shell has been implicated in several aversive behaviors in rats^23,24^. These structural and functional parallels between the OT and NAc, together with evidence that the OT contributes to motivational incentive salience or reward ‘wanting’, raise the possibility that OT might also contain hedonic hotspots capable of enhancing ‘liking’ reactions, similarly to the NAc. However, to our knowledge, no studies have directly tested for hedonic hotspots within the OT.

Here, we used pharmacological microinjections to investigate whether the OTam, previously implicated in reward motivation, contains a hedonic hotspot capable of amplifying ‘liking’ reactions to sucrose taste. Previous studies of the NAc hedonic hotspot of the rostrodorsal medial shell showed that sucrose ‘liking’ reactions were enhanced by local microinjections of any of: the mu opioid receptor agonist DAMGO, orexin-A peptide, or the GABA_A_ receptor agonist muscimol^7,24,25^. Therefore, we tested whether microinjections of any of these three drugs into the anteromedial or anterolateral OT subregions would alter affective ‘liking’ reactions to sucrose taste in rats. Those anatomically distinct OT subregions play different roles in physiological and behavioral responses, and their functional organization is distributed along the medio-lateral axis^26^. Our results indicate that microinjections of DAMGO, orexin, or muscimol into the OTam enhanced hedonic ‘liking’ reactions elicited by oral sucrose infusions over control vehicle levels measured in the same rats, identifying the OTam as an OT hedonic hotspot. In contrast, the same microinjections in the OTal did not enhance ‘liking’ and, in some cases, produced oppositely valenced aversive effects. Furthermore, Fos mapping revealed that DAMGO microinjection into the OTam induced concurrent activation in other hedonic hotspots (the caudal VP and rostromedial OFC), suggesting that the OTam functions as a part of a distributed hedonic hotspot network.

## Materials and Methods

### Animals

Female (n = 20) and male (n = 18) Sprague–Dawley rats (230–460 g) were used. Of these, 14 females and 12 males were assigned to taste reactivity experiments, and 6 females and 6 males to Fos mapping. Rats were housed 2–4 per cage at 21°C under a reverse 12-h light/dark cycle with ad libitum food and water. All procedures were approved by the University of Michigan Institutional Animal Care and Use Committee.

### Surgery

Rats were anesthetized with isoflurane (4–5% induction, 1–2% maintenance) and placed in a stereotaxic apparatus (incisor bar 5.5 mm below intra-aural zero). Bilateral guide cannulas were implanted into either the anteromedial (OTam) or anterolateral (OTal) olfactory tubercle, targeting 2 mm above the dense cell layer. For the OTam-targeted group, skull holes were bilaterally drilled at +2.3 mm anterior and ±3.0 mm lateral to bregma, and the guide cannulas were inserted at a 15° angle toward the midline to a depth of 8.0 mm from the skull surface. For the OTal-targeted group, skull holes were bilaterally drilled at +1.8 mm anterior and ±2.8 mm lateral to bregma, and the guide cannulas were vertically inserted (0° angle) to a depth of 8.0 mm. For taste reactivity experiments, bilateral intraoral cannulas (PE-100 tubing) were implanted to permit infusion of sucrose solutions^27^. Rats received postoperative analgesia and antibiotics and recovered for at least 7 days before testing.

### Drug microinjections

Bilateral microinjections (0.2 μL/side, 1 min) were made using injectors extending 2 mm beyond the guide cannulas. Drugs included the mu-opioid receptor agonist DAMGO (50 ng/side), orexin-A peptide (500 pmol/side), GABAA receptor agonist muscimol (75 ng/side), and ACSF vehicle alone. Each rat received only one treatment per day, and the order was counterbalanced. Drug doses were selected based on previous studies^24,25,28^.

### Taste reactivity tests

The taste reactivity test^2,27,29^ was used to measure affective orofacial reactions elicited by intraoral infusion of a sucrose solution (1% w/v). A 1-mL volume of sucrose solution was delivered into the mouth via oral cannula over a 1-min period. Infusions were administered 25 min after microinjections, approximately when peak pharmacological effects could be expected^24,25,28^. Orofacial and somatic reactions were video-recorded (30 fps) with ventral mirror views. Hedonic ‘liking’ (rhythmic midline tongue protrusions, lateral tongue protrusions, and paw licks, Fig. S1) and aversive ‘disgust’ (gapes, head shakes, face washes, forelimb flails, and chin rubs, Fig. S2) responses were scored offline frame-by-frame using standardized bin-based criteria to ensure equal weighting of frequent and rare behaviors. Totals of hedonic and aversive reactions were computed for each drug and control condition.

### Histology for cannula placement and mapping of behavioral effects

After testing, rats were microinjected with fluorescent tracer (Red RetroBeads) for anatomical verification, then perfused with 4% paraformaldehyde. Brains were sectioned (50 µm) and examined under fluorescence microscopy. Cannula placements were plotted on atlas templates^30^, and functional maps were constructed showing anatomical distributions of hedonic enhancement and suppression relative to vehicle baselines.

### Fos analysis

A separate cohort received DAMGO or vehicle injections in the OTam under identical conditions. Brains were collected 85 min post-injection and processed for Fos immunohistochemistry (rabbit anti-c-Fos (MERCK/Sigma-Aldrich ABE457) and Cy3-conjugated secondary). Local ‘Fos plumes’ around injection sites were quantified as zones of elevated Fos expression relative to vehicle controls, and the average plume radius was used to scale symbol sizes on functional maps (Fig. 5C remapped from Fig. 2A). Fos expression was also quantified in distant mesocorticolimbic structures (OFC, prelimbic cortex, infralimbic cortex, insula, anterior cingulate cortex (ACC), NAc medial shell and core, VP, central amygdala (CeA), basolateral amygdala (BLA), medial amygdala (MeA), lateral hypothalamus (LH), perifornical area of hypothalamus (PFA), arcuate nucleus of hypothalamus, VTA, and paraventricular thalamus) to assess distributed network activation, guided by a template derived from a corresponding brain atlas to ensure consistent placement^31^.

### Statistics

Comparisons of taste reactivity scores were analyzed using repeated-measures one-way ANOVA with Greenhouse–Geisser correction, followed by Dunnett’s post hoc tests to assess drug effects (Fig. 1), or by unpaired t-tests with Welch’s correction to assess regional effects (Figs. 2-4). Comparisons of distant Fos mapping data were analyzed using unpaired t-tests with Welch’s correction (Fig. 5). Statistical significance was set at p < 0.05. All analyses were performed using Graphpad Prism 10 (GraphPad Software, San Diego, CA, USA).

**Figure 1.**
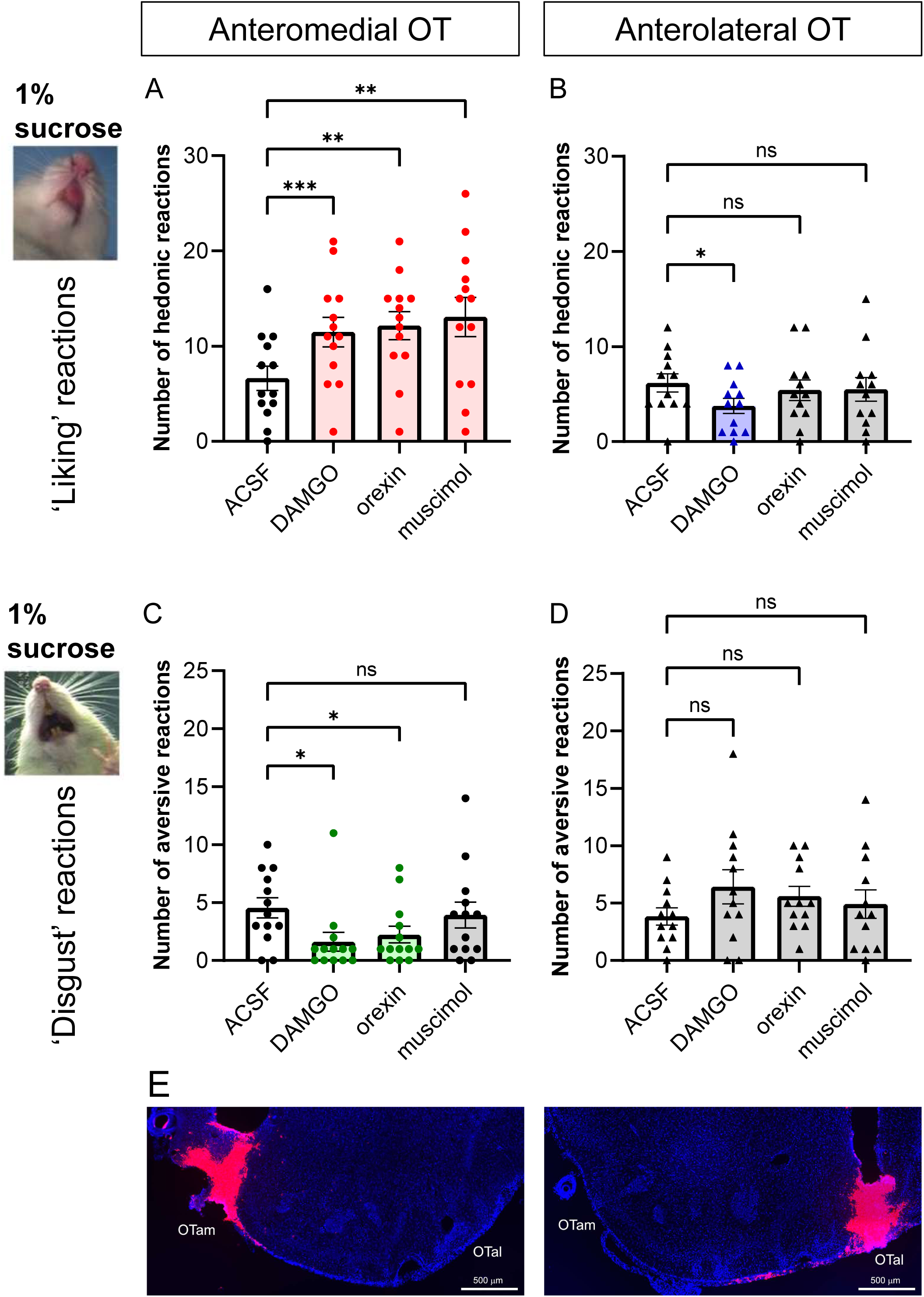
Hedonic ‘liking’ enhancement by microinjections of DAMGO, orexin, and muscimol within the OTam. (A-D) Quantification of hedonic ‘liking’ (A-B) and aversive ‘disgust’ (C-D) reactions to sucrose taste following vehicle (ACSF), DAMGO, orexin, or muscimol microinjections into the OTam (A, C) and (B, D). (E) Representative cannula placements within the OTam (left) and OTal (right) identified by fluorescence tracer (red) and DAPI staining (blue). n = 13 rats for (A, C); n = 12 rats for (B, D). Data are shown as mean ± SEM. *p < 0.05, **p < 0.01 by Dunnett’s test.

**Figure 2.**
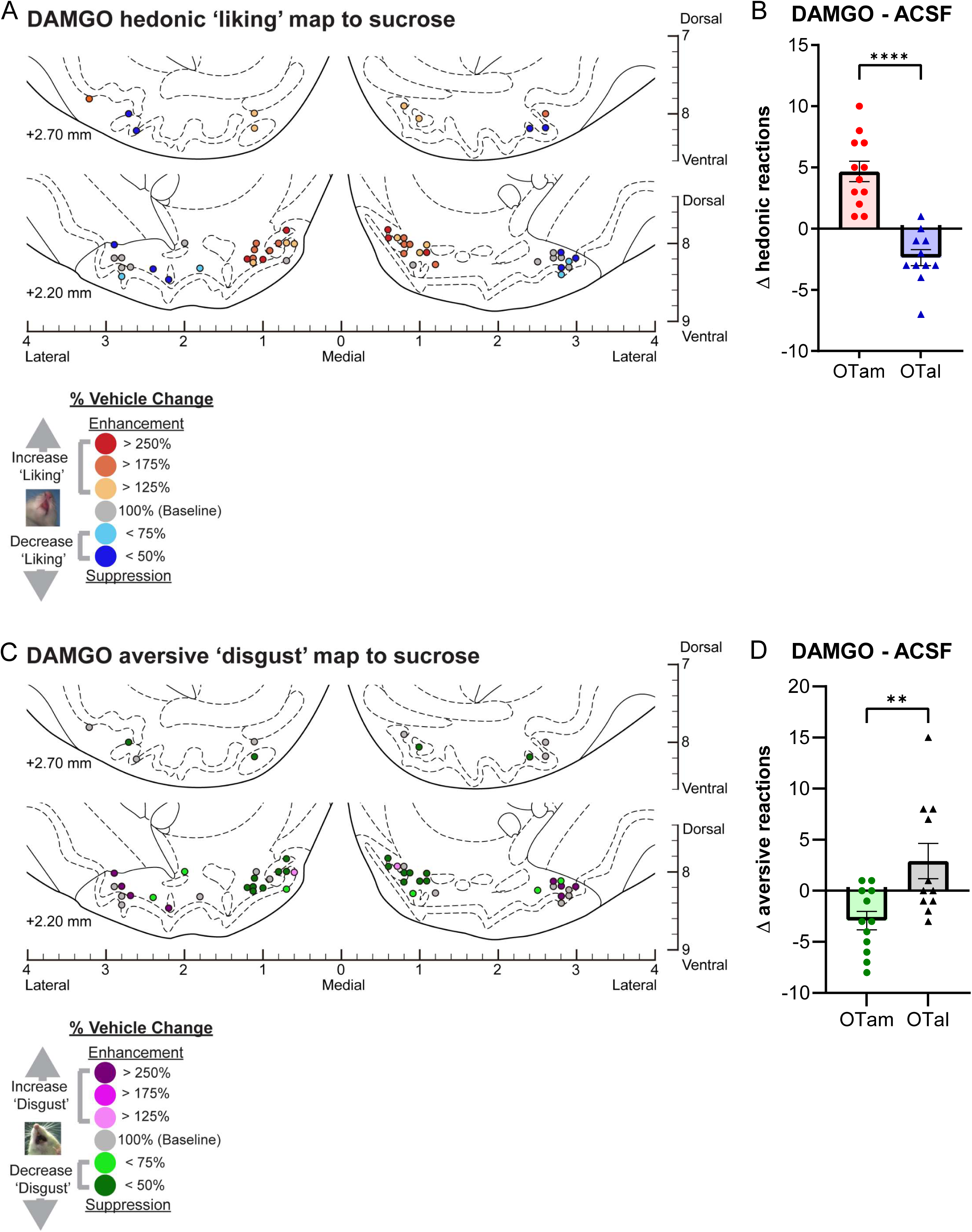
Causation OT maps of hedonic “liking” enhancement by DAMGO. (A) Localization of hedonic function map shows how DAMGO microinjections altered ‘liking’ reactions to intraoral sucrose infusions at each individual OT site. Each symbol indicates the putative cannula tip location for an individual rat plotted on a coronal brain atlas^30^. Symbol color represents within-subject behavioral changes in hedonic reactions induced by DAMGO, expressed as percent change from vehicle ACSF microinjection control levels in the same rat (‘liking’ enhancement: red-yellow; ‘liking’ suppression: blue). (B) DAMGO microinjections differentially alter hedonic ‘liking’ reactions depending on the anatomical subregion of the OT. DAMGO microinjections in the OTam enhanced hedonic ‘liking’ reactions, while those in the OTal suppressed them. (C) Localization of aversive function map shows how DAMGO microinjections altered ‘disgust’ reactions to intraoral sucrose infusions at each individual OT site. Each symbol indicates the putative cannula tip location for an individual rat plotted on a coronal brain atlas^30^. Symbol color represents within-subject behavioral changes in hedonic reactions induced by DAMGO, expressed as percent change from vehicle ACSF microinjection control levels in the same rat (‘disgust’ enhancement: magenta; ‘disgust’ suppression: green). (D) DAMGO microinjections differentially alter aversive ‘disgust’ reactions depending on the anatomical subregion of the OT. DAMGO microinjections in the OTam suppressed aversive ‘disgust’ reactions, whereas those in the OTal showed no significant effect. n = 13 rats for (A-B); n = 12 rats for (C-D). Data are shown as mean ± SEM. ****p < 0.0001; **p < 0.01 by Weltch’s t test.

Further methodological details regarding surgical procedures, drug microinjections, behavioral testing and scoring, and histological analysis are provided in the Supplementary Information.

## Results

### Overview

Microinjections of DAMGO, orexin and muscimol at OTam sites each produced statistically significant enhancements in the number of positive orofacial reactions elicited by the hedonic impact of sucrose taste infusions (Greenhouse–Geisser’s corrected F(1.43, 17.16) = 6.93, p = 0.011; ACSF vs. DAMGO: 95% CI [-6.95, -2.75], p = 0.0001; ACSF vs. orexin: 95% CI [-8.847, -2.230], p = 0.0020; ACSF vs. muscimol: 95% CI [-10.18, -2.746] by Dunnett’s test; Figs. 1A and S1). Sites producing increased hedonic reactions after DAMGO, orexin or muscimol were clustered in close anatomical proximity, revealing the OTam subregion to function as a hedonic hotspot able to enhance sweetness hedonic impact. In contrast, none of the drugs enhanced hedonic reactions when microinjected into OTal sites. Instead, DAMGO microinjection at OTal sites slightly suppressed hedonic reactions to sucrose, indicating a suppressive hedonic coldspot (Greenhouse–Geisser’s corrected F(1.87, 20.59) = 1.72, p = 0.21; ACSF vs. DAMGO 95% CI [0.51 to 4.32], p = 0.014 by Dunnett’s test; Figs. 1B and S1). These results indicate a clear anatomical dissociation between OTam and OTal sites in the effects of DAMGO, orexin and muscimol on hedonic responses to sucrose taste.

Occasionally, rats displayed a few aversive reactions to intraoral infusions of 1% sucrose under the vehicle condition. A repeated-measures one-way ANOVA showed only a trend toward an overall drug effect (Greenhouse–Geisser corrected F(2.34, 28.03) = 3.15, p = 0.051). Nevertheless, post hoc Dunnett’s tests revealed that both DAMGO and orexin microinjections into OTam sites significantly reduced these occasional aversive reactions compared with vehicle (ACSF vs. DAMGO: 95% CI [0.681, 5.17], p = 0.012; ACSF vs. orexin: 95% CI [0.0674, 4.55], p = 0.043), whereas muscimol produced no detectable change (Figs. 1C and S2). No statistically detectable change in aversive responses was observed for any of the three drugs microinjected into the OTal, although there appeared to be a nominal trend towards increased aversion (Greenhouse–Geisser’s corrected F(1.96, 21.6) = 1.55, p = 0.23. Figs. 1D and S2). These results suggest that opioid or orexin stimulation within the OTam hedonic hotspot can both enhance positive hedonic reactions, and suppress negative ‘disgust’ responses.

Cannula placements were confirmed by fluorescent tracer (Red RetroBeads) and DAPI staining (Fig. 1E). In all 13 rats of the OTam-targeted group, cannula tips were localized within the OT, ranging between AP +2.2 to +2.7, ML 0.6 to 1.2, and DV -8.3 to -7.8 mm from bregma. Among the 13 rats of the OTal-targeted group, 12 rats had cannula tips localized within the OT, ranging between AP; +2.2 to +2.7, ML; 1.8 to 3.0, DV; -8.4 to -8.0 mm from bregma; one rat showed a misplaced cannula in the piriform cortex, and its taste reactivity data were excluded from statistical analyses.

### Magnitude of mu-opioid effects on hedonic impact within the OT anteromedial hotspot and anterolateral coldspot

Mu receptor stimulation by DAMGO microinjections within the OTam hotspot nearly doubled the number of hedonic ‘liking’ reactions elicited by sucrose taste (173% of ACSF levels in the same individuals; Fig. 1A). In contrast to these anteromedial enhancements, DAMGO microinjections into the OTal coldspot oppositely decreased hedonic reactions to nearly half of vehicle control levels (61% of ACSF; Fig. 1B). This regional difference between OTam and OTal in the modulation of ‘liking’ reactions to sucrose taste was statistically significant (Welch’s corrected t(20.27) = 6.67, p < 0.001; Figs. 2A-B).

Conversely, DAMGO microinjection within the OTam hedonic hotspot further reduced the relatively rare aversive ‘disgust’ responses to sucrose taste to less than half of vehicle control levels (36% of ACSF; Fig. 1C). DAMGO microinjection within the OTal did not significantly alter the number of ‘disgust’ responses. The regional difference in changes of ‘disgust’ reactions to sucrose taste was again statistically significant (Welch’s corrected t(15.28) = 3.00, p < 0.01; Figs. 2C-D).

### Magnitude of orexin effects within the OT anteromedial hotspot and anterolateral coldspot

Orexin-A peptide microinjections within the OTam hotspot similarly almost doubled the number of hedonic ‘liking’ reactions elicited by sucrose taste (183% of ACSF; Fig. 1A). In contrast, orexin microinjections within the OTal coldspot did not significantly change the number of hedonic reactions elicited by sucrose (Fig. 1B). This regional difference in enhancement of ‘liking’ reactions to sucrose taste was statistically significant (Welch’s corrected t(19.75) = 3.74, p = 0.0013; Figs. 3A-B).

**Figure 3.**
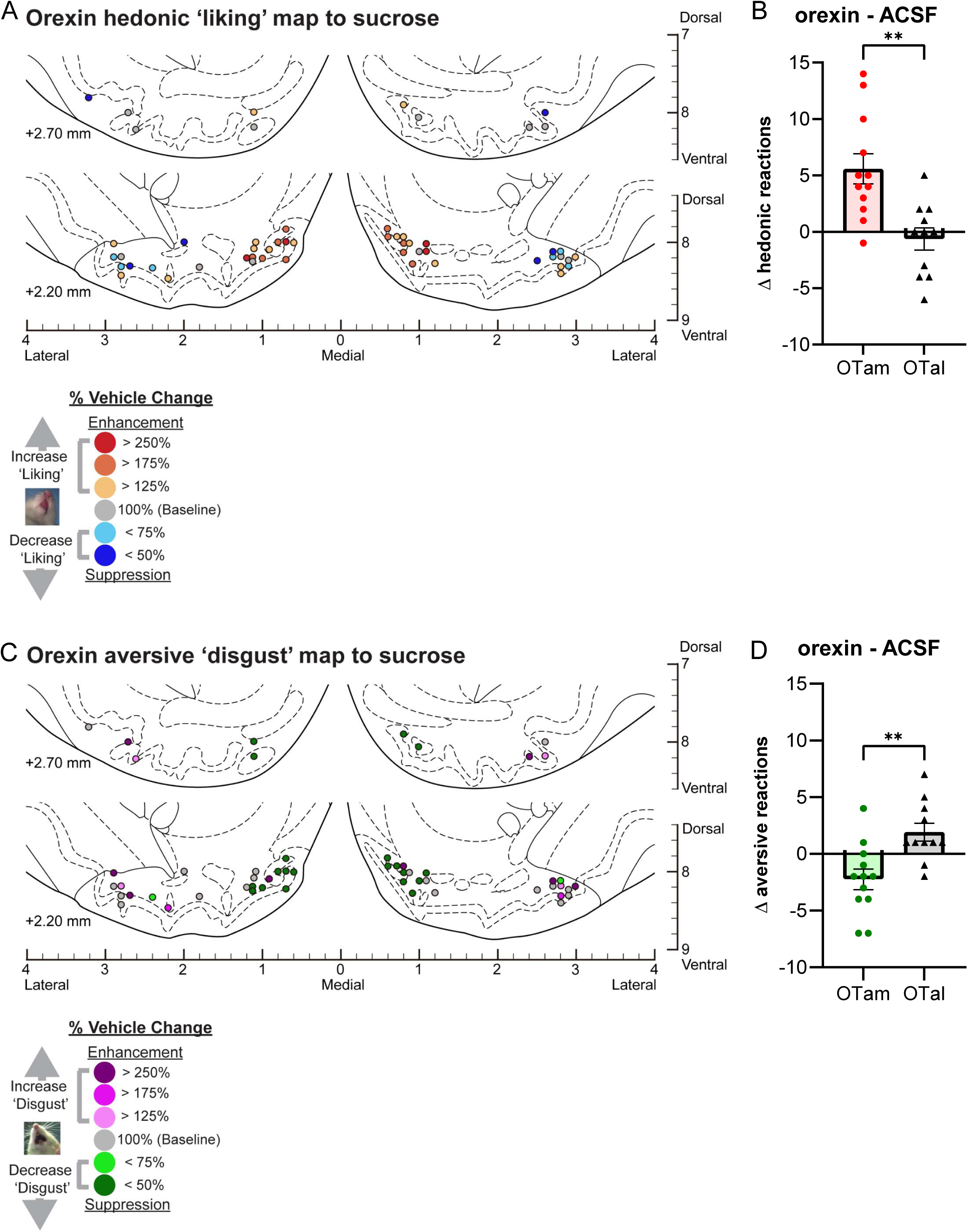
Causation OT maps of “liking” enhancement by orexin. (A) Localization of hedonic function across OT sites showing how orexin microinjections altered ‘liking’ reactions to intraoral sucrose infusions at each individual site. Each symbol represents the putative cannula tip location for an individual rat plotted on a coronal brain atlas^30^. Symbol color indicates within-subject behavioral changes in hedonic reactions induced by orexin, expressed as percent change from vehicle (ACSF) control levels in the same rat (‘liking’ enhancement: red–yellow; ‘liking’ suppression: blue). (B) Orexin microinjections differentially altered hedonic ‘liking’ reactions depending on the anatomical subregion of the OT. Orexin microinjections in the OTam enhanced hedonic ‘liking’ reactions, whereas those in the OTal produced no significant change. (C) Localization of aversive function across OT sites showing how orexin microinjections altered ‘disgust’ reactions to intraoral sucrose infusions at each individual site. Each symbol represents the putative cannula tip location for an individual rat plotted on a coronal brain atlas^30^. Symbol color indicates within-subject behavioral changes in aversive reactions induced by orexin, expressed as percent change from vehicle (ACSF) control levels in the same rat (‘disgust’ enhancement: magenta; ‘disgust’ suppression: green). (D) Orexin microinjections differentially altered aversive ‘disgust’ reactions depending on the anatomical subregion of the OT. Orexin microinjections in the OTam suppressed aversive ‘disgust’ reactions, whereas those in the OTal produced no significant effect. n = 13 rats for (A-B); n = 12 rats for (C-D). Data are shown as mean ± SEM. **p < 0.01 by Welch’s t test.

**Figure 4.**
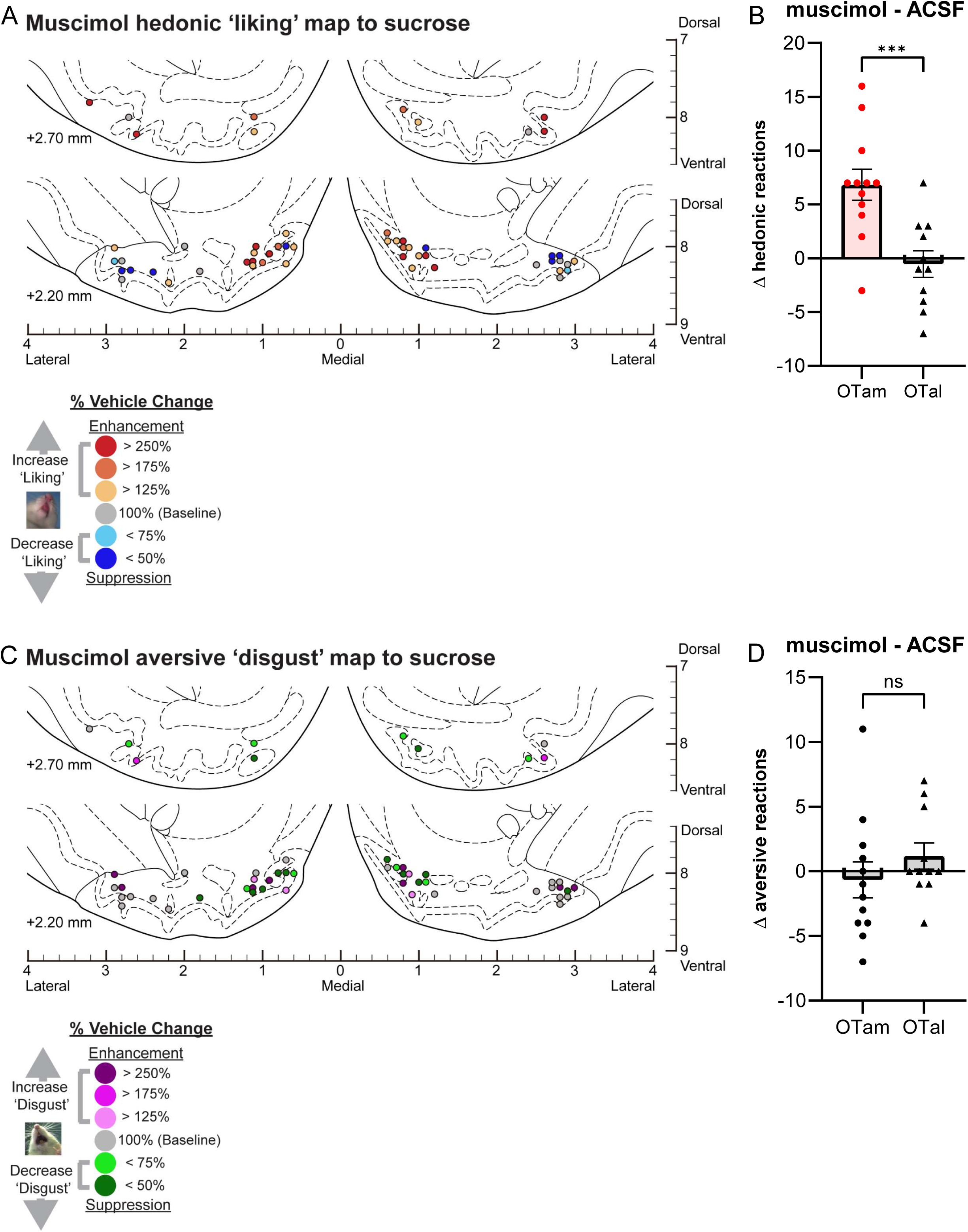
Causation OT maps of “liking” enhancement by muscimol. (A) Localization of hedonic function map shows how muscimol microinjections altered ‘liking’ reactions to intraoral sucrose infusions at each individual OT site. Each symbol indicates the putative cannula tip location for an individual rat plotted on a coronal brain atlas^30^. Symbol color represents within-subject behavioral changes in hedonic reactions induced by muscimol, expressed as percent change from vehicle ACSF microinjection control levels in the same rat (‘liking’ enhancement: red-yellow; ‘liking’ suppression: blue). (B) Muscimol microinjections differentially alter hedonic ‘liking’ reactions depending on the anatomical subregion of the OT. Muscimol microinjections in the OTam enhanced hedonic ‘liking’ reactions, whereas those in the OTal produced no significant change. (C) Localization of aversive function map shows how muscimol microinjections altered ‘disgust’ reactions to intraoral sucrose infusions at each individual OT site. Each symbol indicates the putative cannula tip location for an individual rat plotted on a coronal brain atlas^30^. Symbol color represents within-subject behavioral changes in hedonic reactions induced by muscimol, expressed as percent change from vehicle ACSF microinjection control levels in the same rat (‘disgust’ enhancement: magenta; ‘disgust’ suppression: green). (D) Muscimol microinjections did not significantly alter aversive ‘disgust’ reactions in either the OTam or OTal subregion. n = 13 rats for (A-B); n = 12 rats for (C-D). Data are shown as mean ± SEM. ***p < 0.0001 by Weltch’s t test.

**Figure 5.**
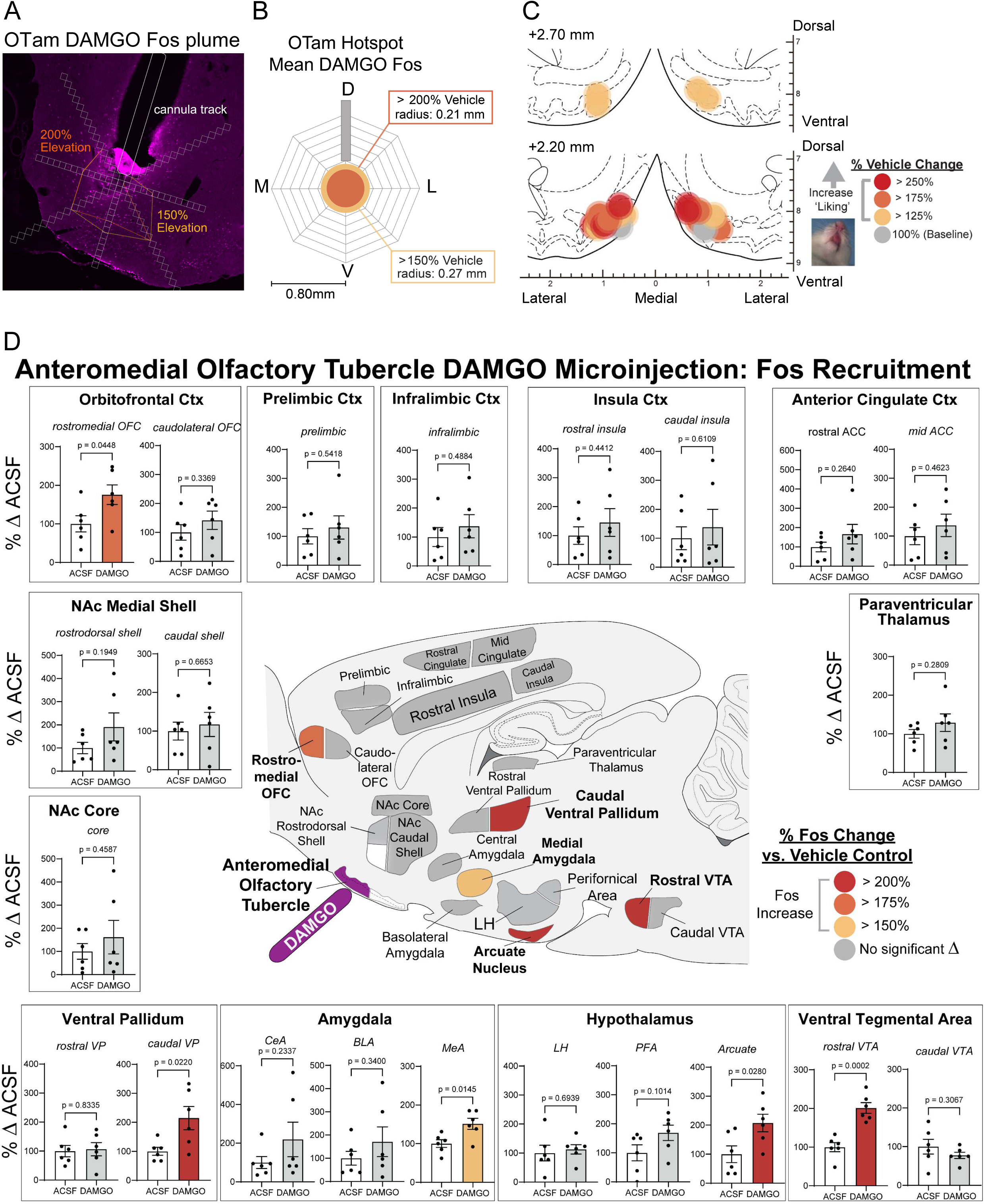
Fos plume and mapping of DAMGO microinjection in the OT anteromedial hotspot. (A) Representative photomicrograph showing local Fos plumes surrounding the cannula tip within the OTam hotspot. (B) Average size of Fos plumes. DAMGO, n = 6 rats; vehicle, n = 6 rats. D: dorsal, M: medial, V: ventral, L: lateral. (C) Remapping of hedonic function localization (from Fig. 2A) within the OTam revealed by DAMGO microinjections, with symbol size reflecting the average radius of Fos plumes. Each symbol represents the putative cannula tip location for an individual rat plotted on a coronal brain atlas^30^. Symbol color indicates within-subject behavioral changes in hedonic reactions induced by DAMGO, expressed as percent change from vehicle (ACSF) control levels in the same rat (‘liking’ enhancement: red–yellow). (D) Distant Fos recruitment following DAMGO microinjection into the OTam hotspot. Brain map illustrates elevated Fos expression in mesocorticolimbic structures recruited for hedonic enhancement. Colors denote significant percent Fos elevation relative to vehicle control rats. Significant Fos increases were observed in other hedonic hotspots, including the caudal VP and rostromedial OFC. Fos elevation was also recruited in other limbic regions, such as rostral VTA, arcuate nucleus of the hypothalamus, and medial amygdala. DAMGO, n = 6 rats; vehicle, n = 6 rats. Bar graphs show mean ± SEM of percent Fos enhancement in each structure relative to vehicle controls. p values are calculated using Weltch’s t test.

Conversely, aversive ‘disgust’ responses to sucrose taste were suppressed to approximately half of vehicle control levels following orexin microinjections within the OTam hotspot (49% of ACSF; Fig. 1C), whereas orexin microinjection within the OTal coldspot did not significantly affect ‘disgust’ responses. This regional difference in the suppression of ‘disgust’ reactions was statistically significant (Welch’s corrected t(20.84) = 3.55, p < 0.01; Figs. 3C-D).

### Magnitude of muscimol effects on hedonic impact within the OT anteromedial hotspot

Muscimol microinjections within the OTam hotspot nearly doubled the number of hedonic ‘liking’ reactions elicited by sucrose taste (198% of ACSF; Fig. 1A). A suppressive coldspot was not detected for muscimol in the OTal, as it did not change the number of hedonic reactions (Fig. 1B). This regional difference between OTam and OTal in the enhancement of ‘liking’ reactions to sucrose taste was statistically significant (Welch’s corrected t(20.75) = 3.87, p = 0.0003; Figs. 4A-B).

The number of aversive ‘disgust’ responses to sucrose taste was not significantly affected by muscimol microinjections in either the OTam hotspot or the OTal coldspot (Figs. 1C-D). Muscimol microinjections therefore did not produce a significant regional difference regarding changes of ‘disgust’ reactions to sucrose taste (Welch’s corrected t(19.68) = 1.07, p = 0.30; Figs. 4C-D).

### Fos plume and mapping: local and distant anatomical spread of drug impact following DAMGO microinjection in the OT anteromedial hotspot

To assess the primary local activation within the anteromedial OT hotspot and its concomitant co-activation in other regions, we conducted Fos plume and distant mapping analyses for DAMGO microinjections, selected as the representative drug. Fos-expressing cells surrounding the cannula tip were quantified to measure local ‘Fos plumes’ (Fig. 5A). These Fos plumes typically exhibited a two-layered structure, consisting of inner zones of intense (>200%) Fos elevation with an average radius of 0.21 mm (volume = 0.039 mm^3^), surrounded by outer zones of moderate (>150%) elevation with an average radius of 0.27 mm (volume = 0.083 mm^3^), relative to baseline Fos levels measured at equivalent sites in vehicle ACSF-injected control rats (Figs. 5A-B). The averaged diameters of these Fos plumes were used to set the symbol sizes in the remapping of the functional hedonic enhancement sites derived from taste reactivity data (Fig. 5C, remapped from Fig. 2A). The color of each symbol represents the intensity of functional effects induced by DAMGO microinjection at that site, expressed as the within-subject percentage change in hedonic ‘liking’ reactions to sucrose compared to vehicle baseline levels measured in the same rat.

We next assessed distant changes in Fos expression across several mesocorticolimbic structures following DAMGO microinjection within the OTam. Fos levels were compared with those of control rats that received vehicle ACSF microinjection at equivalent sites. This analysis aimed to test the hypothesis that stimulation of neurons within a particular hedonic hotspot would recruit Fos elevations in other anatomically distant hedonic hotspots, thereby producing concurrent activation across multiple hotspots as a distributed hedonic network mediating ‘liking’ enhancements. DAMGO microinjection into the OTam hotspot induced 175-200% increases in Fos expression within the caudal VP and the rostromedial OFC hedonic hotspots (Fig. 5D). We did not detect a significant Fos increase in the NAc rostrodorsal medial shell, another known subcortical hedonic hotspot, although some individual rats exhibited marked Fos elevations. Significant Fos increases were also observed in the rostral ventral tegmental area, arcuate nucleus of the hypothalamus, and medial amygdala (Fig. 5D).

## Discussion

### The anteromedial OT as a newly identified hedonic hotspot

Our results demonstrate that the anteromedial OT subregion contains a hedonic hotspot capable of enhancing positively valenced orofacial ‘liking’ reactions to sucrose taste following local mu-opioid, orexin, or GABAergic stimulation. Microinjections of DAMGO, orexin-A, or muscimol within the OTam all significantly increased ‘liking’ reactions to sweetness, similarly as reported for other hedonic hotspots in NAc, VP, etc^4,32^.

In contrast, microinjections of the same drugs into the lateral OT subregion did not enhance ‘liking’ reactions, and DAMGO microinjection instead suppressed hedonic reactions to sweetness. These findings suggest that OTal may serve as a suppressive ‘hedonic coldspot’ able to reduce positive hedonic impact, similar to coldspots previously reported in the caudal NAc medial shell, rostral VP, caudolateral OFC and rostral insula^4,32^. For example, microinjections of mu-, delta-, or kappa-opioid agonists in the caudal NAc medial shell suppress ‘liking’ reactions to sucrose taste^7,28^. Thus, just as the NAc medial shell exhibits a rostro-caudal gradient of hedonic hotspots and coldspots, the OT may display a corresponding medio-lateral gradient, with the OTam acting as a hedonic hotspot and the OTal as a coldspot that each respectively enhance or suppress ‘liking’ reactions to pleasant sucrose taste.

### The OTam hotspot as a component of a distributed hedonic network

Our histochemical analyses confirmed that DAMGO microinjections in OTam produced local ‘Fos plumes’ (0.21-0.28 mm in radius) of neuronal activation surrounding the microinjection site (Fig. 5). This supports the interpretation that DAMGO acted primarily within the OTam itself and did not spread to adjacent OT subregions or the NAc when enhancing ‘liking’ reactions to sucrose taste.

DAMGO microinjection into the OTam also induced distant Fos elevation in other known hedonic hotspots, including the caudal VP and the rostromedial OFC. These findings suggest that the OTam participates in a distributed hedonic network that coordinates pleasure generation across multiple mesocorticolimbic nodes. This is similar to previous reports that opioid microinjections in one hedonic hotspot recruits neurobiological activations in other hedonic hotspots, suggesting the entire network may function as a unitary whole. Further, several additional mesolimbic structures implicated in reward motivation and appetite, including the rostral VTA, medial amygdala, and arcuate nucleus of the hypothalamus were also activated by DAMGO microinjections into the OTam. Further studies are required to delineate the anatomical boundaries of the hedonic hotspot within OTam, or identify the precise circuitry linking the OTam to these downstream structures and to clarify how such network interactions give rise to enhanced hedonic impact.

### Methodological considerations and behavioral limits

In this study, the sucrose concentration was set at a relatively low level (1%) to prevent potential ceiling effects in ‘liking’ responses to overly sweet sucrose that might obscure further neurobiologically-induced hedonic enhancement. To better characterize OTam enhancement characteristics and to confirm the hedonic suppressive effect of the coldspot in OTal, future taste reactivity studies could employ sucrose solutions of appropriately adjusted concentrations.

Conversely, to test whether neurochemical stimulation of the OTam hedonic hotspot can reduce negatively valenced ‘disgust’ reactions, and whether stimulation of the OTal hedonic coldspot can amplify them, future studies could use bitter quinine solutions^7^. In the present study, only a very few ‘disgust’ gapes or related aversive reactions were observed in response to 1% sucrose, as expected, indicating that a more unpleasant bitter stimulus may be required to evaluate modulation of negatively valenced aversive reactions.

### Neurochemical and circuit mechanisms: opioid, orexin, and GABAergic systems

Our data suggest that mu receptor systems in the OTam play a causal role in enhancing ‘liking’ reactions to sweet sucrose. The endogenous opioid system involves β-endorphin-mu receptor, dynorphin-kappa receptor and enkephalin-delta receptor pairs. Notably, members of all three peptide families can activate each receptor subtype to varying degrees^33^. The OT express mu, kappa, and delta receptors similar to the NAc^34–37^. In addition, MSNs in the OT express preproenkephalin and prodynorphin genes^15^. In the NAc hedonic hotspot, ‘liking’ reactions are enhanced by microinjections of a mu agonist, a delta agonist, or even a kappa agonist^28^, and future studies should compare the effects of delta and kappa stimulation in the OTam hotspot. We speculate that activation of the OTam by sensory stimuli may trigger the release of endogenous OT enkephalin and possibly dynorphin under certain conditions, which in turn could amplify hedonic impact^38^.

The OTam also shows higher expression of orexin-1 receptor (Hcrtr1) than the lateral OT^39^. This differential expression of orexin receptor may partly underlie the enhanced hedonic ‘liking’ reactions produced by orexin microinjection within the OTam. Local microinjection of an orexin antagonist into the mouse OTam reportedly induces aversive behavioral responses to food associated-odors that are otherwise attractive^40^, suggesting that orexin signaling within the OTam contributes to assigning positive valence to odor cues. Orexin microinjections into hedonic hotspots of NAc, VP, OFC, and insula can increase ‘liking’ reactions to sucrose taste similarly to opioid microinjections^5,25,41^. Further investigation is needed to determine whether orexin signals in the OTam hotspot enhance ‘liking’ reactions, and might be modulated by changes in physiological states such as those accompanying hunger, satiety, or circadian rhythms.

Muscimol microinjection within the OTam hotspot also enhanced hedonic reactions to sucrose taste, consistent with previous reports of GABA agonist effects within the NAc hedonic hotspot^24^. A major hypothesis of NAc reward function posits that relative inhibition of NAc MSNs mediates reward functions by reducing tonic MSN GABAergic suppression of downstream targets in the VP, VTA, and lateral hypothalamus, thereby releasing those reward-related targets into relative excitation^42–48^. Whether a similar mechanism operates in the OT has remained unknown, but our muscimol results are consistent with the hypothesis that GABAergic inhibition of MSNs in amOT disinhibits downstream targets into relative excitation to cause hedonic enhancement. Among possible targets, both D1- and D2-expressing OT MSNs appear to send axons toward or through the VP, and D1-expressing OT MSNs may form direct synaptic connection with the VTA^12,49,50^. Further understanding of the synaptic connectivity and the inhibitory-excitatory dynamics within the OTam will be essential to explain how local GABAergic inhibition by muscimol can ultimately enhance hedonic impact.

### Implications for olfaction, flavor, and food pleasure

As its olfactory label implies, the OT receives dense synaptic inputs from the olfactory bulb and olfactory cortex, which likely convey odor information to the OT. Olfaction plays important roles in flavor perception through the retronasal pathway, an essential contributor to the pleasure of eating^51,52^. The importance of olfaction in food pleasure is highlighted by clinical findings that patients with olfactory dysfunctions often experience a marked reduction in eating enjoyment^53,54^.

As a speculative hypothesis, the hedonic hotspot identified in the OTam may contribute to the pleasurable impact of flavor sensations, thereby promoting learning and motivation toward particular foods. Supporting this idea, human neuroimaging studies have been shown that subjective pleasantness ratings of odors correlate with activity in the OT^55^. Our identification of a hedonic hotspot within the OTam thus provides new insight into affective neuroscience by revealing a neural substrate that potentially links olfaction, flavor and the pleasure of eating.

## Acknowledgements

We thank Ileana Morales, Yan Xiong, David Nguyen, Katie Emery, Madeliene Granillo, Aishwarya Ramaswami, Carina Castellanos, Yugo Fukazawa, Kazuki Kuroda, Eri Murai, Takako Maegawa, and Mayumi Yamamoto for their generous support.

## Author Contributions

KM designed the study, conducted and analyzed the behavioral experiments and histology, and wrote the manuscript. KCB contributed to study design, interpretation of the data, and revision of the manuscript.

## Supplementary Materials and Methods

### Surgical procedures

Rats were anesthetized with isoflurane gas prior to surgery (induction: 4–5%, maintenance: 1–2%) and placed in a stereotaxic apparatus (David Kopf Instruments) with the incisor bar set 5.5 mm below the intra-aural zero. At the onset of surgery, atropine (0.05 mg/kg, i.p.), cefazolin (75 mg/kg, s.c.), and carprofen (5 mg/kg, s.c.) were administered.

Permanent microinjection guide cannulas (9 mm, 26-gauge C315G or 23-gauge C317G; Plastics One) were bilaterally implanted into the OT, targeting either the OTam (n = 13 for the taste reactivity test; n = 12 for Fos mapping) or the OTal (n = 13 for the taste reactivity test). The cannula tips were positioned approximately 2 mm above the dense cell layer of the OT.

For the OTam-targeted group, skull holes were bilaterally drilled at +2.3 mm anterior and ±3.0 mm lateral to bregma, and the guide cannulas were inserted at a 15° angle toward the midline to a depth of 8.0 mm from the skull surface. For the OTal-targeted group, skull holes were bilaterally drilled at +1.8 mm anterior and ±2.8 mm lateral to bregma, and the guide cannulas were vertically inserted (0° angle) to a depth of 8.0 mm. Cannula coordinates were made as bilaterally symmetrical as possible across individuals within each target group. Guide cannulas were anchored to the skull with surgical screws and dental acrylic. Dummy cannulas (33-gauge C315DCS or 30-gauge C317DCS; Plastics One) were inserted and kept in place at all times, except during behavioral testing, to prevent occlusion.

For rats in the taste reactivity group, bilateral intraoral cannulas (polyethylene PE-100 tubing) were implanted during the same surgery to permit oral infusions of sucrose solutions. Oral cannulas entered the oral cavity at the upper cheek pouch lateral to the first maxillary molar, ascended beneath the zygomatic arch, and exited the skin at the dorsal head cap^27^. The cannulas did not disrupt normal feeding behavior.

Postoperatively, rats received carprofen at 24 and 48 h and cefazolin at 24 h after surgery, and were allowed to recover for one week before behavioral testing.

### Drug microinjections

Rats were gently cradled by hand on the experimenter’s lap during microinjections. Polyethylene PE-20 tubing was connected to microinjection cannulas (33-gauge C315I or 30-gauge C317I, Plastics One) that extended 2 mm beyond the guide cannulas to reach OT targets. Drug or vehicle (artificial cerebrospinal fluid, ACSF) solutions were brought to room temperature (∼21°C), and inspected to confirm the absence of precipitation before microinjection.

Drugs and vehicle solutions were freshly prepared at the beginning of each test series and stored frozen across consecutive test days. All drugs were dissolved in ACSF and bilaterally microinjected over a 1-min period at a volume of 0.2 μl per side (0.2 μl/min) by syringe pump. Injectors were left in place for 1 min following microinjection to allow diffusion, after which dummy cannulas were replaced, and rats were immediately placed in the taste reactivity testing chamber.

Four microinjection solutions were tested in each rat: DAMGO, a mu receptor agonist (50 ng/0.2 μl per side); orexin-A peptide (500 pmol/0.2 μl per side); muscimol, a GABA_A_ receptor agonist (75 ng/0.2 μl per side), and ACSF vehicle alone (0.2 μl per side, vehicle control). Drug doses were selected based on previous studies^24,25,28^.

For taste reactivity tests, each rat received bilateral microinjections of only one drug or vehicle per day. The order of drugs and vehicle was counterbalanced among DAMGO, orexin, and ACSF for the first three daily microinjections across rats, with muscimol being tested on the fourth day in all rats.

### Taste reactivity tests

The taste reactivity test^2,27,29^ was used to measure affective orofacial reactions elicited by intraoral infusion of a sucrose solution (1% w/v). A 1-mL volume of sucrose solution was delivered into the mouth via oral cannula over a 1-min period. Infusions were administered 25 min after microinjections, approximately when peak pharmacological effects could be expected^24,25,28^. To infuse sucrose solution into the mouth, a syringe containing sucrose solution was mounted on a syringe pump and connected via polyethylene tubing (PE-50 attached to a PE-10 delivery nozzle) to the rat’s oral cannula. Orofacial and somatic reactions were video-recorded at 30 frame-per-second using a close-up lens and an angled mirror placed underneath the transparent floor to capture ventral views for subsequent video analysis.

Prior to testing, rats were extensively handled to familiarize them with the experimenters. They were then habituated to the test chamber for 25 min on three consecutive days and received a mock microinjection of vehicle ACSF on the third day of habituation.

### Taste reactivity video scoring

Hedonic, aversive, and neutral taste reactivity patterns were scored off-line using frame-by-frame video analysis. Hedonic responses were defined as rhythmic midline tongue protrusions, lateral tongue protrusions, and paw licks^1^. Aversive responses were defined as gapes, head shakes, face washes, forelimb flails, and chin rubs. Neutral responses included passive dripping of the solution from the mouth, ordinary grooming, and rhythmic mouth movements.

A time-bin scoring procedure was employed to ensure that taste reactivity components of different relative frequencies still contributed equally to the final totals, so that frequent components such as rhythmic tongue protrusions did not swamp rare but equally informative components, such as lateral tongue protrusions^1^. Specifically, rhythmic mouth movements, passive dripping, and paw licking reactions, which occur in long bouts, were scored in 5-s time bins (e.g., 5 s of continuous paw licking behavior was counted as one bout). Rhythmic midline tongue protrusions and chin rubs, which occur in shorter bouts, were scored in 2-s bins. Lateral tongue protrusions, gapes, forelimb flails, and head shakes, which typically occur as discrete events, were scored as single occurrences each time they occurred (e.g., one gape equals one occurrence).

Individual totals were calculated separately for hedonic and aversive categories. The hedonic reaction total was quantified as the sum of lateral tongue protrusion, rhythmic tongue protrusion, and paw lick scores. The aversive reaction total was quantified as the sum of gape, head shake, face wash, forelimb flail, and chin rub scores.

### Histology for cannula placement and mapping of behavioral effects

Following behavioral testing, rats received microinjection of a fluorescent tracer (Red RetroBeads; 0.2 μL per side) through the microinjection cannulas to allow subsequent anatomical mapping of injection sites. Rats were then deeply anesthetized with an overdose of sodium pentobarbital, and perfused transcardially with 0.1M sodium phosphate buffer (NaPB), followed by 4% paraformaldehyde. Brains were removed, post-fixed in 4% paraformaldehyde for 1 day, and then immersed in 25% sucrose (in 0.1 M NaPB) for 1-2 days. Brains were frozen, coronally sectioned at 50 μm on a freezing microtome, mounted, air-dried, and coverslipped using DAPI-containing mounting medium (ProLong Gold Antifade Mountant with DAPI). Bilateral microinjection sites for each rat were assessed by fluorescence microscope, and plotted onto corresponding coronal maps adapted from a rat brain atlas^30^. These plots were used to extrapolate the locations of individual injection sites onto composite group maps of the OT to identify functional hedonic hotspots and coldspots.

Group maps of the behavioral effects of OT microinjections were constructed in the coronal plane to display all injection sites on the same map, revealing functional differences across the mediolateral extents of the anterior OT. Functional effects on hedonic reactions were visualized using color-coding to represent the percentage change in affective behaviors for each rat, calculated as the number of hedonic or aversive reactions during drug conditions divided by the number of corresponding reactions elicited by sucrose under the vehicle ACSF condition (Figs. 2-4). To avoid division by zero for rats that showed no hedonic or aversive reactions under ACSF, we added +1 to the number of reactions in all microinjection conditions (ACSF, DAMGO, orexin, and muscimol) for those rats before calculating percentages. Map symbols indicate the putative tip position of the injection cannulas, as verified by fluorescent tracer labeling.

### Histological Fos analysis following DAMGO microinjection in the OTam hotspot

Fos analysis was conducted in a separate group of rats from those used in taste reactivity tests to reveal the anatomical spread of neural activation caused by drug microinjections. Distant Fos effects were also examined in the same rats to determine whether DAMGO in the OTam recruited neuronal activity in other mesocorticolimbic structures, including other hedonic hotspots such as in orbitofrontal cortex, ventral pallidum, etc. Rats in the Fos analysis group received DAMGO or vehicle ACSF microinjection in the OTam under conditions identical to those of the behavioral group on their first test day. Cannula placements in the Fos group corresponded to those in the taste reactivity test group.

### Immunohistochemistry

Eighty-five minutes after microinjection, rats were deeply anesthetized with a lethal dose of sodium pentobarbital (150-200 mg/kg) and transcardially perfused with 0.1M NaPB followed by 4% paraformaldehyde. Brains were removed, post-fixed in 4% paraformaldehyde for 1 day, and transferred to 25% sucrose solution (in 0.1 M NaPB) for 1-2 days. Coronal sections (25 μm) were cut on a cryostat (Leica) and processed for Fos immunohistochemistry.

Sections were rinsed three times for 10 min in 0.1 M NaPB, blocked in 5% normal donkey serum / 0.2% Triton-X in PBS for 60 min, and incubated overnight with a polyclonal rabbit anti-c-Fos primary antibody (1:1000; MERCK/Sigma-Aldrich ABE457). After three 10-min rinses in 0.1M NaPB, sections were incubated for 2-h with a Cy3-conjugated donkey anti-rabbit secondary antibody (1:500; Jackson ImmunoResearch). Sections were rinsed again, mounted, air-dried, and coverslipped with a DAPI-containing mounting medium (ProLong Gold Antifade Mountant with DAPI). Fos immunoreactivity was imaged using a fluorescence slide scanner (VS200, Evident Scientific), and images of whole-brain coronal sections were captured at 20x magnification using Olyvia software.

### Local Fos Plume Analysis

Fos plumes were mapped at 20x magnification by counting Fos-positive cells within consecutive 50 µm x 50 µm tissue blocks along seven radial arms emanating from the center of the microinjection site (45°, 90°, 135°, 180°, 225°, 270°, and 315°). Counting proceeded outward along each arm until at least two consecutive blocks contained no Fos-positive cells. Percent increases in DAMGO-induced Fos expression were calculated relative to baseline levels measured at the same sites in vehicle-injected control rats (Fig. 5A). Fos plumes were defined as zones of intense elevation (>200%) or moderate elevation (>150%), and radii were averaged across the seven arms to determine plume size. The averaged plume radii were used to scale symbol sizes in functional maps of hedonic enhancement sites (Fig. 5C). Cannula tip locations were plotted onto corresponding atlas maps to construct OTam functional localization figures (Fig. 5C, remapped from Fig. 2A).

### Distant Fos Mapping Analysis

Functional activation of circuitry recruited by DAMGO microinjection into the OTam hotspot immediately prior to euthanasia was assessed by quantifying Fos expression at distant sites across multiple structures: OFC, prelimbic cortex, infralimbic cortex, insula, anterior cingulate cortex (ACC), NAc medial shell and core, VP, central amygdala (CeA), basolateral amygdala (BLA), medial amygdala (MeA), lateral hypothalamus (LH), perifornical area of hypothalamus (PFA), arcuate nucleus of hypothalamus, VTA, and paraventricular thalamus.

Within each subregion, Fos-positive cells were counted in two to six sample boxes (200 µm x 200 µm), placed equidistantly within the structure, and approximately matched across rats, guided by a template derived from a corresponding brain atlas to ensure consistent placement^31^. The number of sample boxes was adjusted for each structure to capture an average 6–15 Fos-positive cells in vehicle-injected control rats. Counts across sample boxes were summed to determine total Fos expression for each subregion or structure.

**Supplementary Figure S1.**
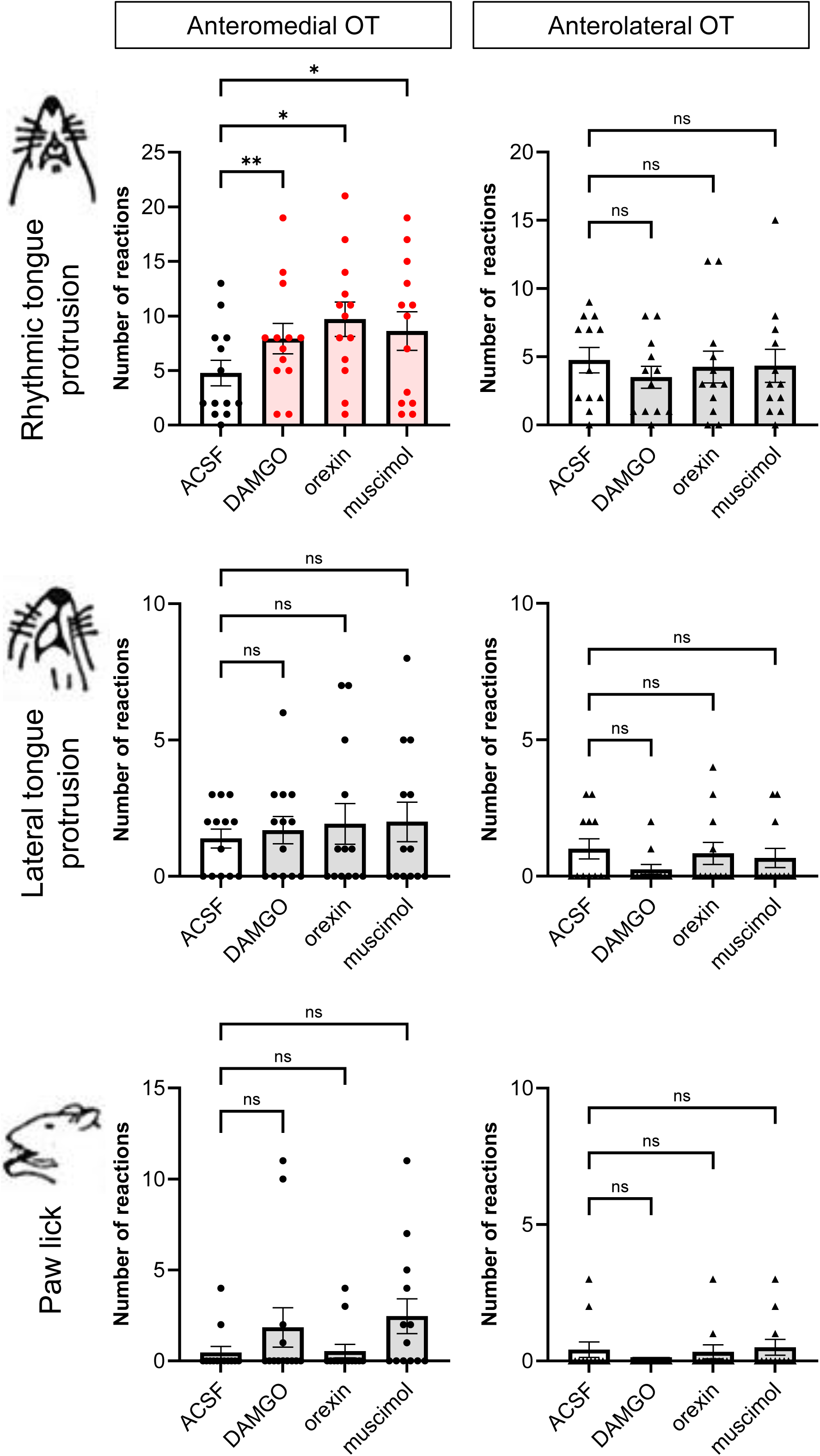
Breakdown of hedonic ‘liking’ reactions. Fig. S1 decomposes the total hedonic ‘liking’ reaction scores shown in Fig. 1 A,B into their constituent affective facial expressions. Left panels depict rats receiving drug microinjections into the anteromedial OT, and right panels depict injections into the anterolateral OT. The three rows show the positive ‘liking’ components scored during sucrose tasting: top, rhythmic tongue protrusions; middle, lateral tongue protrusions; bottom, paw licks. For each drug condition, values represent the mean number of reactions elicited during the taste reactivity test. Summing these three affective components yields the total ‘liking’ scores presented in Fig. 1A,B. *p < 0.05, ns not significant. n = 13 rats for anteromedial OT; n = 12 rats for anterolateral OT.

**Supplementary Figure S2.**
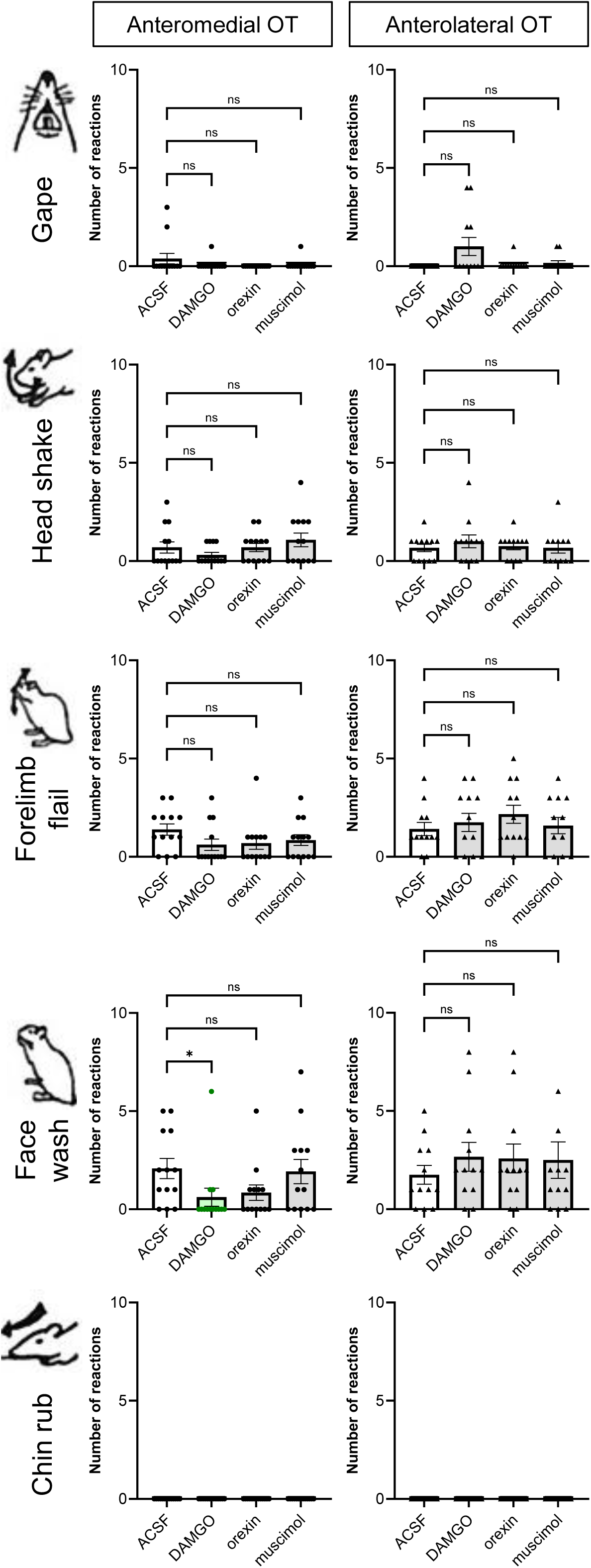
Breakdown of aversive ‘disgust’ reactions. Fig. S2 decomposes the total aversive ‘disgust’ reaction scores shown in Fig. 1C,D into their constituent negative affective facial and somatic expressions. Left panels depict rats receiving drug microinjections into the anteromedial OT, and right panels depict injections into the anterolateral OT. The five rows illustrate the aversive ‘disgust’ components scored during sucrose tasting under drug conditions: top, gapes; second, head shakes; third, forelimb flails; fourth, face washes; bottom, chin rubs. For each drug condition, values represent the mean number of reactions elicited during the taste evaluation period. Summing these five aversive components yields the total ‘disgust’ scores presented in Fig. 1C,D. *p < 0.05, ns not significant. n = 13 rats for anteromedial OT; n = 12 rats for anterolateral OT.

